# SignaFish: a zebrafish-specific signaling pathway resource

**DOI:** 10.1101/041822

**Authors:** Kitti Csályi, Dávid Fazekas, Tamás Kadlecsik, Dénes Türei, Leila Gul, Balázs Horváth, Dezső Módos, Amanda Demeter, Nóra Pápai, Katalin Lenti, Péter Csermely, Tibor Vellai, Tamás Korcsmáros, Máté Varga

## Abstract

Understanding living systems requires an in depth knowledge of the signaling networks that drive cellular homeostasis, regulate intercellular communication and contribute to cell fates during development. Several resources exist to provide high-throughput datasets or manually curated interaction information from human or invertebrate model organisms. We previously developed SignaLink, a uniformly curated, multi-layered signaling resource containing information for human and for the model organisms nematode *Caenorhabditis elegans* and fruit fly *Drosophila melanogaster*. Until now, the use of the SignaLink database for zebrafish pathway analysis was limited. To overcome this limitation we created SignaFish (http://signafish.org), a fish-specific signaling resource, built using the concept of SignaLink. SignaFish contains more than 200 curation based signaling interactions, 132 further interactions listed in other resources, and it also lists potential miRNA based regulatory connections for 7 major signaling pathways. From the SignaFish website, users can reach other web resources, such as ZFIN. SignaFish provides signaling or signaling-related interactions that can be examined for each gene, or downloaded for each signaling pathway. We believe that the SignaFish resource will serve as a novel navigating point for experimental design and evaluation for the zebrafish community and for researchers focusing on non-model fish species, such as cyclids.

---

The biology of living organisms cannot be interpreted without an in depth knowledge of the signaling networks that drive cellular homeostasis, regulate intercellular communication and contribute to establish cell fates, which have an indispensable role both during and after development. While early research concentrated on deciphering individual signaling pathways, later results demonstrated that these pathways are often unexpectedly interwoven [1, 2]. This particular feature of signaling networks can often contribute to seemingly confusing results upon the analysis of interactions between signaling proteins. Without accounting for the details of an interaction (direction or sign), the interpretation of experimental data will be often erroneous.

A better understanding of signaling network topology allows for a better accounting of the observed phenotypes by recognizing robustness within the examined network or identifying key proteins, which could become potential targets in future treatments or interventions [3]. Hence the need for well annotated protein-protein interaction (PPI) and signaling databases for biological and biomedical research.

SignaLink (http://signalink.org/) [4, 5], a uniformly curated, multi-layered signaling resource has become over the years a widely used signaling resource for the model organisms nematode *Caenorhabditis elegans* and fruit fly *Drosophila melanogaster*, as well as a gap-filling human database with high coverage. With an easy-to-use interface, customizable download site and integrated datasets, it had been used in several high profile studies (for example [6]) and also integrated in model-organism resources, such as FlyBase and WormBase.

However, the use of SignaLink for the zebrafish community was limited. To overcome this limitation, we developed SignaFish (http://signafish.org), a fish-specific signaling resource, built using the concept of SignaLink.

With SignaFish, we have created a manually curated database of seven major signaling pathways, which are biochemically and evolutionary defined, and encompass all major developmental signaling mechanisms [1]: Rtk (receptor tyrosine kinase), Tgf-β (transforming growth factor beta), Wnt/Wingless, Hedgehog, Jak/Stat, Notch and NHR (Nuclear hormone receptor). The pathway interactions were coming from (1) review-based literature searches, which identified the primary zebrafish-related paper that described the interactions; or (2) we used the signaling interactions of *C. elegans, D. melanogaster* and *H. sapiens* from SignaLink2 [5], and based on the interolog and signalog concepts [7, 8], we predicted potential zebrafish signaling interactions. We included a predicted interaction to the SignaFish database if (1) we found no direct evidence that the given interaction has been verified in zebrafish, (2) the orthologs of potentially interacting proteins were present and interacted in at least one species in SignaLink, (3) and, to minimize the false positive interactions, a zebrafish study showed that genetic modification of both potentially interacting proteins produces a phenotype related to the annotated pathway.

SignaFish contains altogether 217 signaling interactions found with manual curation or with high-confident predictions (as described above). These interactions were extended with protein-protein interactions (imported from [9, 10]), and potential miRNA-mRNA connections (from TargetScanFish [11]). Altogether, SignaFish contains pathway and interaction information for 389 proteins and 178 miRNAs (Figure 1). The number of manually curated interactions directly found in zebrafish studies is low (86) compared to the total number of papers available for zebrafish, because we applied a strict curation protocol [4], which allowed the inclusion of only those studies where the identified interaction was investigated between two zebrafish proteins, and the experiments were carried out in zebrafish. Due to the lack of zebrafish specific regulatory resources listing transcription factor – target gene connections, we were not able to include transcriptional regulations to this version of SignaFish. The database structure of SignaFish already allows the incorporation of such transcriptional data upon a large dataset will be available.

**Figure 1.**
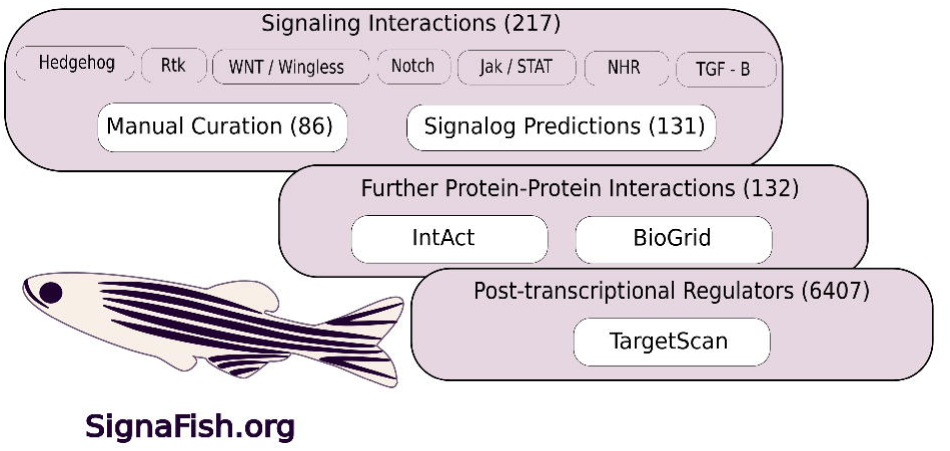
The structure of SignaFish and its sources. The number of interactions acquired from each sources is shown in parenthesis. All parts of the SignaFish database can be browsed and downloaded through the website, where users can filter to specific signaling pathways.

To facilitate the exploration of this complex resource, we implemented it to a website with a user-friendly graphical interface, available at http://signafish.org. The website is linked to main web resources including UniProt, ENSEMBL and ZFIN.

We clearly recognize that the list of the included data is far from complete and we are committed in updating SignaFish annually. We welcome inputs from the wider community of zebrafish researchers, therefore, we included an online form (available from the website) that could be used to submit novel interactions, which – after manual verification – will be included in the next update of SignaFish.

Since its publication the SignaLink database has proven its usefulness for invertebrate model organism, therefore we believe SignaFish will also become an important resource for zebrafish research. The signaling interactions that can be examined for each gene, or downloaded for each signaling pathway will serve as a novel navigating point for experimental design and evaluation for the zebrafish community and for researchers focusing on non-model fish species.

## Acknowledgement

We thank the technical help of Kata Ferenc. This work was supported by research grants from the Hungarian Science Foundation [grant numbers: OTKA K115378, K109349, K115378, NK78012], by the Institute of Advanced Studies Kőszeg (iASK, Kőszeg, Hungary), a Technology Innovation Fund grant from the Hungarian National Developmental Agency (KTIA-AIK-2012-12-1-0010). TD was supported by the EMBL Interdisciplinary Postdoc Programme (EIPOD) under Marie Skłodowska-Curie COFUND Actions (Grant number 291772). TK is a Computational Biology Fellow at The Genome Analysis Centre in partnership with the Institute of Food Research (Norwich, UK), and strategically supported by Biotechnological and Biosciences Research Council, UK. MV was supported by János Bolyai and MedInProt fellowships of the Hungarian Academy of Sciences.

